# Reading canonical and modified nucleotides in 16S ribosomal RNA using nanopore direct RNA sequencing

**DOI:** 10.1101/132274

**Authors:** Andrew M. Smith, Miten Jain, Logan Mulroney, Daniel R. Garalde, Mark Akeson

**Author notes:** Correspondence should be addressed to M.A.

## Abstract

The ribosome small subunit is expressed in all living cells. It performs numerous essential functions during translation, including formation of the initiation complex and proofreading of base-pairs between mRNA codons and tRNA anticodons. The core constituent of the small ribosomal subunit is a ∼1.5 kb RNA strand in prokaryotes (16S rRNA) and a homologous ∼1.8 kb RNA strand in eukaryotes (18S rRNA). Traditional sequencing-by-synthesis (SBS) of rRNA genes or rRNA cDNA copies has achieved wide use as a ‘molecular chronometer’ for phylogenetic studies ^1^, and as a tool for identifying infectious organisms in the clinic ^2^. However, epigenetic modifications on rRNA are erased by SBS methods. Here we describe direct MinION nanopore sequencing of individual, full-length 16S rRNA absent reverse transcription or amplification. As little as 5 picograms (∼10 attomole) of E. coli 16S rRNA was detected in 4.5 micrograms of total human RNA. Nanopore ionic current traces that deviated from canonical patterns revealed conserved 16S rRNA base modifications, and a 7-methylguanosine modification that confers aminoglycoside resistance to some pathological E. coli strains. This direct RNA sequencing technology has promise for rapid identification of microbes in the environment and in patient samples.

---

Nanopore-based direct RNA strand sequencing ^3^ is conceptually similar to nanopore DNA sequencing. An applied voltage across a single protein pore in an impermeable membrane results in an ionic current through the pore ^4^. This current varies when a DNA or RNA strand is captured by the electric field and then moved through the pore in single nucleotide steps regulated by a processive enzyme ^5–7^. The output is a time series of discrete ionic current segments that correspond to the sequence of bases that occupy the pore at any given time ^8,9^. Other PCR-free RNA sequencing technologies (often referred to as direct RNA sequencing because the RNA is present during sequencing) have been implemented using SBS combined with optical readout of fluorophore-labelled DNA nucleotides ^10,11^. They share some of the benefits of nanopore direct RNA sequencing (e.g. absence of PCR biases), however their reported read lengths are short (typically <25 nt ^10^ and <34 nt ^11^ respectively).

Direct nanopore RNA sequencing was first implemented by Oxford Nanopore Technologies (ONT) for mRNA using adapters designed to capture polyadenylated RNA strands ^3^. We reasoned that this technique could be modified to sequence 16S rRNA. 16S rRNA is a logical substrate for nanopore sequencing because of its abundance and broad use for identifying bacteria and archaea. In addition, numerous antibiotics target prokaryotic ribosomes ^12^ which can acquire resistance via nucleotide substitutions, or by gain or loss of base modifications ^13^. These base modifications are difficult to detect using indirect SBS methods. A significant advantage of nanopore sequencing is that modifications can be resolved because each nucleoside touches the nanoscale sensor as the strand translocates through the pore.

Figure 1a illustrates the strategy we used to prepare 16S rRNA for MinION sequencing. Briefly, 16S rRNA was ligated to an adapter bearing a 20-nt overhang complementary to the 3′-end of the 16S rRNA (**Figure 1a** and **Supplemental Fig. 1a**). This overhang included the Shine-Dalgarno sequence ^14^, which targets the conserved anti-Shine-Dalgarno sequence in prokaryotic 16S rRNA ^15^. Next, a modular Oxford Nanopore Technologies (ONT) adapter bearing a proprietary RNA motor protein was hybridized and ligated to the adapted RNA strands thus facilitating capture and sequencing on the MinION.

**Figure 1.**
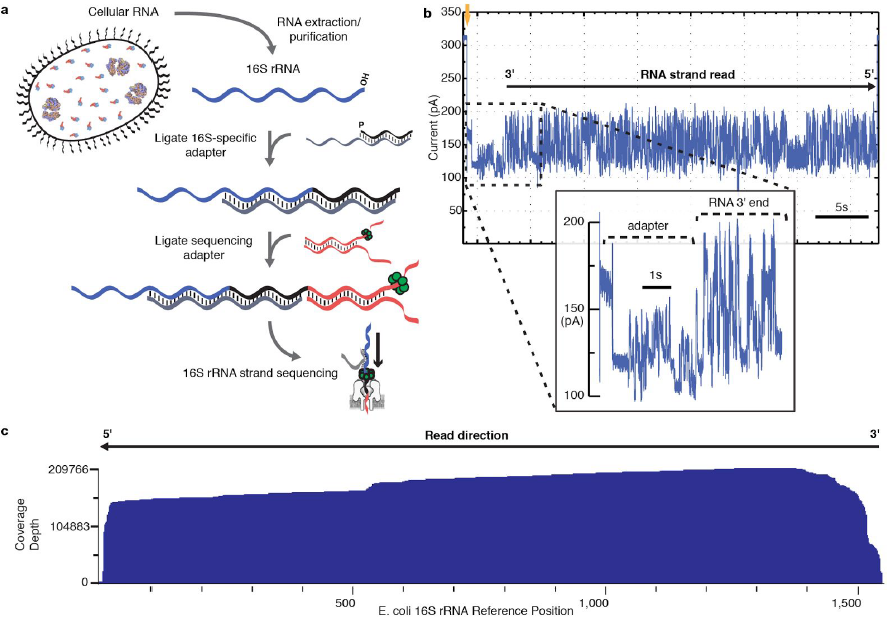
Direct nanopore sequencing of individual *E. coli* 16S ribosomal RNA strands. **(a)** Library preparation for MinION sequencing. Following RNA extraction, a 16S rRNA-specific adapter is hybridized and ligated to the 16S rRNA 3′ end. Next, a sequencing adapter bearing a RNA motor protein is hybridized and ligated to the 3′ overhang of the 16S rRNA adapter. The sample is then loaded into the MinION flowcell for sequencing. **(b)** Representative ionic current trace during translocation of a 16S rRNA strand from *E. coli* str. MRE600 through a nanopore. Upon capture of the 3′ end of an adapted 16S rRNA, the ionic current transitions from open channel (310 pA; gold arrow) to a series of discrete segments characteristic of the adapters (inset). This is followed by ionic current segments corresponding to base-by-base translocation of the 16S rRNA. The trace is representative of thousands of reads collected for individual 16S rRNA strands from *E. coli*. **(c)** Alignment of 200,000+ 16S rRNA reads to *E. coli* str MRE600 16S rRNA *rrnD* gene reference sequence. Reads are aligned in 5′ to 3′ orientation, after being reversed by the base-calling software. Numbering is according to canonical *E. coli* 16S sequence. Coverage across reference is plotted as a smoothed curve. In this experiment, 94.6% of reads that passed quality filters aligned to the reference sequence. Data presented here are from a single flow cell.

Figure 1b shows a representative ionic current trace caused by translocation of a purified *E. coli* 16S rRNA strand through a nanopore in the MinION array. The read begins with an ionic current pattern characteristic of the ONT RNA sequencing adapter strand followed by the 16S rRNA adapter strand. The 16S rRNA is then processed through the nanopore one base at a time in the 3′ to 5′ direction. The ionic current features are typical of long nucleic acid polymers processed through a nanopore ^16,17^.

Sequencing of purified 16S rRNA from *E. coli* strain MRE600 produced 219,917 reads over 24 hours that aligned to the reference sequence (16S rRNA *rrnD* gene) (**Figure 1c**). This represents 94.6% of the total MinION read output for that experiment. Median read length was 1349 bases. We identified 142,295 reads that had sequence coverage within twenty-five nucleotides of the 16S rRNA 5′-end and within fifty nucleotides of the 3′-end.

We calculated the percent read identities for sequence data from 16S rRNA and Enolase 2 RNA (a calibration standard supplied by ONT) (**Supplemental Fig. S2**). The median read identity for 16S rRNA was 81.6% compared to 87.1% for Enolase 2 (**Supplemental Table S1**). Close examination of 16S rRNA reads revealed frequent deletion errors in G-rich regions, which are abundant in non-coding structural RNAs such as 16S rRNA (**Supplemental Table S2 and Supplemental Fig. S3**). This is observed as drops in coverage when unsmoothed read coverage is plotted across the *E. coli* 16S rRNA reference (**Supplemental Fig. S2**). Other sequencing errors may represent true single nucleotide variants (SNVs) from the 16S rRNA reference sequence used for alignment. *E. coli* strains typically have seven 16S rRNA gene copies, with some of the gene copies differing by as much as 1.1%. Modified nucleotides could also alter ionic current from canonical nucleotides ^18,19^. *E. coli* 16S rRNA contains 12 known nucleotide modifications ^20^.

We predicted that both SNVs and nucleoside modifications would result in reproducible nanopore base-call errors. Therefore, we looked for positions that were consistently mis-called relative to the *E. coli* MRE600 16S rRNA reference. Using marginCaller at a posterior probability threshold of 0.3 ^16^, we detected 24 such positions in the nanopore 16S rRNA reads (**Supplemental Table S3**). Five of these were mis-calls resulting from minor variants in the reference sequence relative to the other 16S rRNA gene copies. For example, at position 79 the reference is adenine (A79), whereas the other six 16S rRNA gene copies have a guanosine, in agreement with the majority of nanopore reads. One of the highest probability variants was at G527 in the reference, which was systematically mis-called as a C (**Figure 2a & b**). This residue is located in a conserved region of the 16S rRNA 530 loop, near the A-site in the ribosome ^21^. The guanosine base at this position is known to be methylated at N7 (m7G527) ^22^, which creates a delocalized positive charge. We hypothesized that this modification would significantly alter the ionic current segments that contain m7G527, thus resulting in the systematic base-call error.

**Figure 2.**
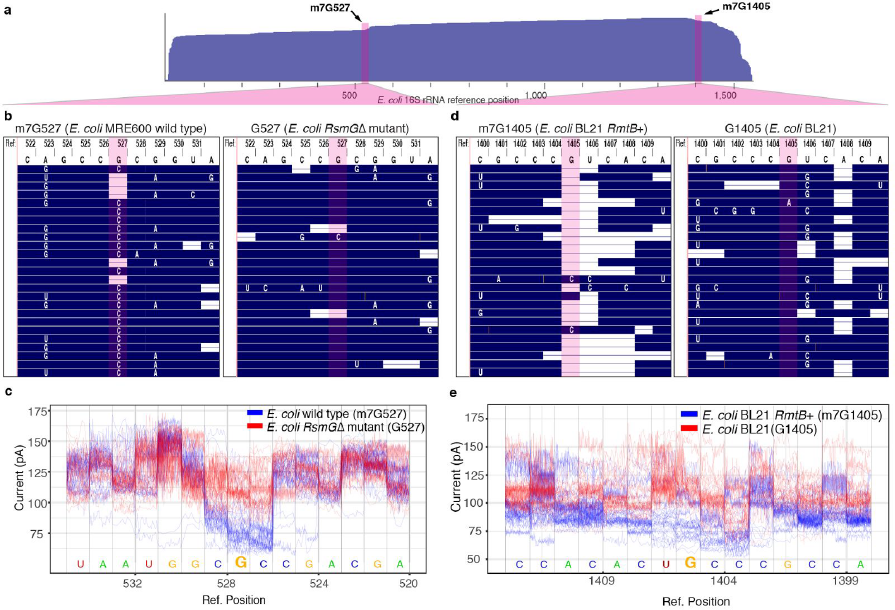
Detection of 7mG modifications in *E. coli* 16S rRNA. **(a)** Diagram showing the positions along *E. coli* 16S rRNA that correspond to the expanded sequence alignments in panels b-e. Arrows indicate the positions of G527 and G1405 in the *E. coli* reference. **(b)** Alignment of nanopore RNA sequence reads proximal to position 527 of *E. coli* 16S rRNA. Numbered letters at the top represent DNA bases in the reference 16S rRNA gene. Blue regions in the body of the panel denote agreement between reference DNA bases and nanopore RNA strand base-calls. White letters denote base call differences between the reference and the nanopore reads, and horizontal white bars represent base deletions in the nanopore RNA reads. Columns highlighted in red correspond to position 527. The left inset is *E. coli* str. MRE600 (wild type) 16S rRNA (m7G527), and the right inset is *RsmG* mutant strain 16S rRNA (canonical G527). **(c)** Nanopore ionic current traces proximal to position 527 of the *E. coli* 16S rRNA reference. Blue traces are for wild type *E. coli* 16S rRNA translocation events bearing m7G at position 527. Red traces are for mutant strain 16S rRNA translocation events bearing a canonical G at position 527. **(d)** Alignment of nanopore RNA sequence reads proximal to position 1405 of *E. coli* 16S rRNA. Use of colors, shapes, and letters are as described for panel (b). The left inset is engineered mutant *E. coli* str. BL21 (*RmtB*+) 16S rRNA (m7G1405); the right inset is *E. coli* str. BL21 16S rRNA (G1405). **(e)** Nanopore ionic current traces proximal to position 1405 of the *E. coli* 16S rRNA reference. Blue traces are for mutant strain 16S rRNA translocation events bearing m7G at position 1405. Red traces are for wild type 16S rRNA translocation events bearing a canonical G at position 1405.

To test this hypothesis, we compared *E. coli* str. MRE600 (wild type) 16S rRNA nanopore reads with reads for an *E. coli* strain that lacks the enzyme RsmG, which is responsible for N7 methylation at G527 ^23^. We validated the absence of methylation at G527 in the *RsmG* deficient strain by chemical cleavage (**Supplemental Fig. S4a**). As predicted, a canonical guanosine base at position 527 in the mutant strain eliminated the reproducible base-call error seen in the wild type *E. coli* strain (**Figure 2b**). Examination of ionic current segments containing G527 and m7G527 in RNA strands for the respective strains confirmed that m7G alters ionic current relative to canonical G (**Figure 2c**).

Typically, *E. coli* 16S rRNA contains only one m7G at position 527. However, some pathogenic strains that are resistant to aminoglycosides contain an additional m7G at position 1405 ^24^. The enzymes responsible for G1405 methylation, such as RmtB ^25^, are thought to have originated from microbes that produce aminoglycosides and are shuttled on multidrug-resistance plasmids ^26^. Given the pronounced signal difference for m7G at position 527, we thought it should also be possible to detect m7G in this context.

To this end, we engineered an *E. coli* strain that carried *RmtB* on an inducible plasmid (pLM1-RmtB, see Methods). We confirmed that this *RmtB*+ strain was aminoglycoside resistant, (**Supplemental Fig. S4b**) consistent with N7 methylation of G1405. We then compared 16S rRNA sequence reads for this strain (*RmtB*+) with reads from the parent *E. coli* strain (BL21) without the plasmid (**Figure 2a & d**). We observed an increase in deletions and base mis-calls in 16S rRNA reads for the *RmtB*+ strain at position G1405 and the adjacent U1406. These mis-calls were absent in the 16S rRNA reads for the parent BL21 strain, which bears a canonical guanosine at G1405. Examination of ionic current segments containing G1405 and m7G1405 in RNA strands for the respective strains confirmed that m7G alters ionic current relative to canonical G (**Figure 2e**), as was observed at position 527. In this region, methylated cytosines at positions 1402 and 1407 may also contribute to the aberrant ionic current, which could account for the base mis-calls proximal to those bases in the parent strain (**Figure 2d,** right panel).

Nanopore detection of epigenetic RNA modifications is not limited to m7G. While examining base mis-calls proximal to G527, we also noted a systematic miscall at U516 (**Supplemental Fig. S5**). This mis-called position had the highest probability variant in our marginCaller analysis (**Supplementary Table S3)**. We hypothesized that this was due to pseudouridylation at U516 which is typical in *E. coli* 16S rRNA ^27^. As a test, we compared nanopore reads for the wild type strain with reads for a mutant strain (*RsuA*Δ) bearing a canonical uridine at position 516. We found that mis-calls and ionic current deviations present at U516 in the wild type were absent in the mutant strain (**Supplemental Fig. S5**) consistent with the hypothesis.

Another important feature of direct nanopore 16S rRNA reads is that they are predominantly full-length. It has been established that more complete 16S rRNA sequences allow for improved taxonomic classification ^28^. To test if full-length MinION 16S rRNA reads gave better classification than short reads, we sequenced purified 16S rRNA from three additional microbes (*Methanococcus maripaludis* str. S2, *Vibrio cholerae* str. A1552, and *Salmonella enterica* str. LT2). These were chosen to give a range of 16S rRNA sequence similarities to *E. coli* (68.1%, 90.4%, and 97.0% identity respectively). The 16S rRNA adapter sequence was altered slightly for each microbe (see Methods). We binned reads by length, sampled 10,000 reads per bin for each microorganism, mixed them *in silico*, and aligned them to 16S rRNA sequences for all four microbes. A read was counted as correctly classified if it aligned to a 16S rRNA reference sequence for the source microorganism. As predicted, the classification accuracy increased with read length from 67.9% for short reads (200-600 bases) to 96.9% for long reads (>1000 bases) (**Figure 3a**). When using all the reads for each bin per microbe (i.e. no sampling), the average classification accuracy increased to 97.8% for long reads (>1000 bases) (**Supplemental Fig. S6**).

**Figure 3.**
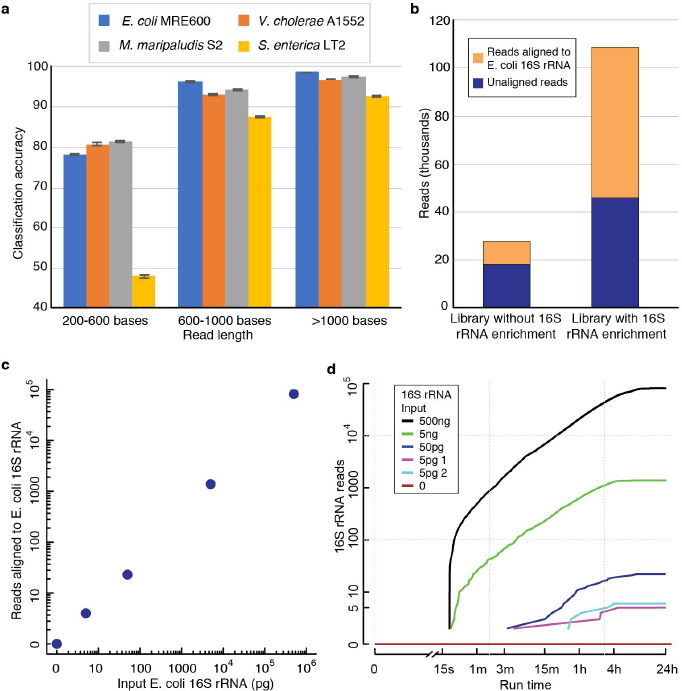
Direct 16S rRNA sequencing discriminates among microbes and can detect *E. coli* 16S rRNA at low concentration in a human RNA background. **(a)** Classification accuracy from an *in silico* mixture of 16S rRNA reads from four microbes. Reads were binned based on length and 10 iterations of classification using 10,000 randomly sampled reads per microbe were performed. A read was called as correctly classified if it aligned to one of the 16S rRNA reference sequences for that microbe. The error bars indicate one standard deviation for the 10 iterations. **(b)** 16S rRNA sequencing yield for libraries prepared from *E. coli* str. K12 total RNA with and without enrichment. Sequencing libraries were prepared from 5 µg total RNA. The enrichment library used a desthiobiotinylated version of the 16S rRNA-specific adapter, which was hybridized and selected for using magnetic streptavidin beads (see Methods). The two 16S rRNA sequencing libraries were then prepared essentially the same way. **(c)** 16S rRNA reads from sequencing libraries prepared with *E. coli* str. MRE600 16S rRNA titered into 4.5 µg total RNA from HEK 293T cells. **(d)** 16S read accumulation over time in titration sequencing runs. The lines correspond to libraries shown in **c**.

Our early sequencing experiments required purifying 16S rRNA, which is prohibitively slow for clinical applications. Therefore, we devised an enrichment strategy that permits selective preparation of 16S rRNA from total bacterial RNA. This involved adding a desthiobiotin to the 16S rRNA adapter (see Methods). The adapter was hybridized to 16S rRNA in a mixture, and then bound to streptavidin-conjugated magnetic beads. This allowed washing and removal of non-specific RNA. The library preparation was then performed as usual. To test the enrichment method, we prepared 16S rRNA sequencing libraries from the same *E. coli* total RNA preparation with and without the enrichment step. We observed that enrichment increased the number of reads that aligned to 16S *E. coli* rRNA sequence >5-fold relative to the library without enrichment (**Figure 3b**).

This suggested that 16S rRNA could be selectively sequenced from a human total RNA background, at relative proportions that would be expected in a clinical sample. To test this, we titered 5 pg to 500 ng of *E. coli* 16S rRNA into 4.5 µg total RNA from human embryonic kidney cells (HEK 293T) and prepared sequencing libraries (**Figure 3c**). The lowest mass (5 pg) approximates the amount of 16S rRNA from 300 *E. coli* cells ^29^. 4.5 µg of total human RNA approximates the total RNA typically extracted from 1 ml of blood.

We observed a linear correlation between *E. coli* 16S rRNA reads and *E. coli* 16S rRNA concentrations over a 100,000-fold sample range (**Figure 3c)**. In replicate 5 pg experiments, we observed only 4-5 16S rRNA reads, which nonetheless could be distinguished from the total human RNA negative control (0 16S rRNA reads in 24 hours). Because nanopore data are collected in real-time, we examined how rapidly *E. coli* 16S rRNA was detected in these MinION runs. We extracted acquisition times for all reads that aligned to *E. coli* 16S rRNA (**Figure 3d**). At concentrations ≥5ng, we found that the first 16S rRNA read occurred within ∼20 seconds of the start of sequencing. This means that some 16S rRNA strands were immediately captured and processed by the MinION upon initiation of the sequencing run. At lower input amounts (<5 ng), we detected *E. coli* 16S rRNA strands in less than one hour. Combined with library preparation, this suggests that nanopore sequencing could detect microbial 16S rRNA in a complex clinical or environmental sample within 2 hours.

Some nanopore RNA sequencing applications (e.g. strain-level taxonomic identification or detection of splice sites in transcript isoforms) will require better base-call accuracy than achieved in this study. These improvements seem likely based on prior evidence for MinION DNA sequencing where base call accuracies increased from 66% in 2014 ^16^ to 92% in 2015 ^30^.

It is plausible that nanopore RNA sequencing will work for all classes of RNA, with long reads providing more complete sequence and modification annotations.

## Acknowledgements

Harry Noller and Hossein Amiri provided purified *E. coli* 16S rRNA. Laura Lancaster advised on primer extension experiments. Manny Ares provided assistance with *S. enterica* cell culture and advised on 16S rRNA methylation experiments. David Alexander provided HEK 293T cells. Fitnat Yildiz and Jennifer Teschler provided *V. cholerae* total rRNA. Todd Lowe provided *M. maripaludis* total RNA. Oxford Nanopore Technologies provided direct RNA sequencing kits and Enolase 2 mRNA. Ariah Mackie proofread the final draft. Discussions with James Hadfield reaffirmed our interest in sequencing 16S rRNA. Brad Bebout gave useful feedback on the potential utility of 16S rRNA nanopore sequencing in the field. This work was supported by NHGRI grant HG006321 (MA).

## Author contributions

AMS designed and performed RNA bench experiments, conceived and designed MinION experiments, helped perform MinION experiments and bioinformatics, and co-wrote the paper. MJ helped conceive and design MinION experiments, helped perform MinION experiments and bioinformatics, and co-wrote the paper. LM helped design *RmtB* experiments and engineered an *E. coli* strain that carried *RmtB*. DRG co-wrote the manuscript and helped conceive and design MinION experiments. MA co-wrote the manuscript, helped conceive and design experiments, and oversaw the project.

## Materials and Methods

### Cell culture and total RNA Isolation for 16S rRNA sequencing

*E. coli* strains BW25113 JW3718Δ and BW25113 JW2171Δ (strains hereafter referred to by gene deletion names *RsmG*Δ and *RsuA*Δ, respectively), deficient for 16S rRNA modifying enzymes RsmG and RsuA respectively, were purchased from the Keio Knockout collection ^31^ (GE Dharmacon). *E. coli* strains K12 MG1655, *RsmG*Δ, *RsuA*Δ and *S. enterica* strain LT2 were grown in LB media (supplemented with 50 µg/ml kanamycin for *RsmG*Δ and *RsuA*Δ) at 37°C to an A_600_ = 0.8-1.0. Cells were harvested by centrifugation and total RNA was extracted with Trizol (Thermo Fisher) following the manufacturer’s recommended protocol. All total RNA samples were treated with DNase I (NEB) (2U/10 ug RNA) in the manufacturer’s recommended buffer at 37°C for 15 minutes. Following the DNase I reaction, RNA was extracted by acid phenol/chloroform extraction (pH 4.4, Fisher Scientific) and two rounds of chloroform extraction. RNA was precipitated with sodium acetate (pH 5.2) and ethanol. RNA was resuspended in nuclease-free water and stored at −80°C. For experiments where human RNA was used as a background, total RNA was extracted from 10^7^ HEK 293T cells following the same steps.

### 16S rRNA purification

*E. coli* strain MRE600 16S rRNA was isolated from sucrose-gradient purified 30S subunits. *Vibrio cholerae* strain A1552 and *Methanococcus maripaludis* strain S2 16S rRNAs were isolated by gel purification from total RNA. 50-100 µg total RNA (DNase I treated) was heated to 95°C for 3 minutes in 7M urea/1xTE loading buffer and run on a 4% acrylamide/7M urea/TBE gel for 2.5 hours at 28W. Gel bands corresponding to 16S rRNA were cut from the gel. 16S rRNA was electroeluted into Maxi-size D-tube dialyzers (3.5 kDa MWCO, EMD Millipore) in 1X TBE for 2 hours at 100V. RNA was precipitated with sodium acetate and ethanol overnight at −20°C. RNA was pelleted washed once with 80% ethanol. Recovered RNA was resuspended in nuclease free water and quantitated using a Nanodrop spectrophotometer.

### Oligonucleotides and 16S rRNA adapters

The 16S rRNA adapter was designed as a double-stranded DNA oligo. The bottom 40-nt strand has one 20-nucleotide region complementary to the 3’ end of the 16S RNA, and a second 20-nt region complementary to the top strand (**Supplemental Fig. S1a**), with the sequence 5′-CCTAAGAGCAAGAAGAAGCCTAAGGAGGTGATCCAACCGC-3′. The top strand, which is directly ligated to the 16S rRNA, used the sequence 5′-pGGCTTCTTCTTGCTCTTAGGTAGTAGGTTC-3′ (p, 5′ phosphate). For *V. cholerae* and *M. maripaludis*, the 3′ terminal 20-nt of the bottom strand were slightly changed to yield adapters with perfectly complementary to their respective 16S rRNA 3′ ends. This resulted in the strands 5′-CCTAAGAGCAAGAAGAAGCCTAAGGAGGTGATCCAGCGCC-3′ and 5′-CCTAAGAGCAAGAAGAAGCCAGGAGGTGATCCAGCCGCAG-3′, respectively. To make a 16S rRNA adapter, top and the bottom strands were hybridized at 10 µM each in a buffer containing 10 mM Tris-HCl (pH 8.0), 1 mM EDTA, and 50 mM NaCl. The mixtures were heated to 75°C for 1 minute before being slowly cooled to room temperature in a thermocycler. We confirmed the adapter hybridizes and ligates to *E. coli* str. MRE600 16S rRNA 3′ end by a gel electrophoresis-based assay with a 6-FAM-labeled version of the top strand (**Supplemental. Fig. S1b**). For experiments where 16S rRNA was enriched from a total RNA background, a desthiobiotin was added to the 5′ terminus of the bottom strand. All adapter oligonucleotides were synthesized by IDT.

### Purified 16S rRNA Sequencing Library Preparation

Sequencing libraries of purified 16S rRNA for *E. coli* str. MRE600, *V. cholerae* str. A1552, and *M. maripaludis* str. S2 were prepared as follows: 2 pmol 16S rRNA adapter and 1.5 µg purified 16S rRNA (approximately 3 pmol) were added to a 15 µL reaction in 1x Quick Ligase buffer with 3000U T4 DNA ligase (New England Biolabs). The reaction was incubated at room temperature for 10 minutes. These reactions were cleaned up using 1.8x volume of RNAclean XP beads (Beckman Coulter), washed once with 80% ethanol and resuspended in 20 µl nuclease-free water. The RNA sequencing adapter (Oxford Nanopore Technologies) was ligated to the RNA library following manufacturer recommended protocol.

### Preparation of RNA Sequencing libraries enriched for 16S rRNA

Enrichment-based 16S sequencing libraries were prepared for *E. coli* strains K-12 MG1655, BL21 DE3 pLys, BL21 DE3 pLys pLM1-*RmtB+*, BL21 DE3 pLys pLM1-*RmtBΔ*, *RsmG*Δ, *RsuA*Δ, and *S. enterica* strain LT2. 16S rRNA-enriched sequencing libraries were essentially prepared as described for purified 16S rRNA with the following exceptions: 15 pmol of 5′ desthiobiotinylated 16S rRNA adapter was added to 4.5-5 µg total RNA in 10 µL buffer containing 10 mM Tris-HCl pH 8, 1 mM EDTA and 50 mM NaCl. The mixture was heated to 50°C for 1 minute and slowly cooled to room temperature in a thermocycler (∼10 minutes). The mixture was then incubated at room temperature for 20 minutes with 100 µL MyOne C1 magnetic streptavidin beads (Thermo Fisher) in 10 mM Tris-HCl (pH 8), 1 mM EDTA, 500 mM NaCl, and 0.025% NP-40 (Buffer A). The beads were washed once with an equal volume of Buffer A and once with an equal volume of buffer containing 10 mM Tris-HCl (pH 8), 1 mM EDTA, 150 mM NaCl (Buffer B). To elute 16S rRNA-enriched RNA, 20 µl Buffer B amended with 5 mM biotin was incubated with the beads at 37°C for 30 minutes. The hybridized 16S rRNA adapter was then ligated by bringing the mixture to 40 µL 1x Quick Ligase buffer (New England Biolabs) and adding 3000U of T4 DNA ligase (New England Biolabs). The rest of the library preparation was performed the same as described for purified 16S rRNA sequencing libraries.

### *In vivo* methylation of 16S rRNA G1405

The *RmtB* gene was purchased as a synthetic gBlock from IDT with the sequence from GenBank accession EU213261.1. pET-32a+ (EMD Millipore) and *RmtB* gBlock were digested with XhoI and NdeI. Digested plasmid and gBlock were ligated with T4 DNA ligase (NEB) to create plasmid pLM1-*RmtB*+. To create *RmtB* null plasmid, pLM1-*RmtB*Δ, XhoI and NdeI digested pET-32a+ was end repaired and ligated. Plasmids were transformed into *E. coli* DH5a cells (NEB) and confirmed by Sanger sequencing. Confirmed clones for pLM1-*RmtB*+ and pLM1-*RmtB*Δ were transformed into *E. coli* BL21 DE3 pLysS cells to create expression strains. To methylate G1405 in 16S rRNA, *E. coli* BL21 DE3 pLys pLM1-*RmtB*+ cells were cultured in 150 ml LB at 37°C with Ampicillin (100 ug/ml) until OD_600_ ∼ 0.4. Cultures were diluted into 1 L in pre-warmed LB media with Ampicillin (100 ug/ml), and plasmid expression was induced with 1 mM IPTG. Cultures were grown at 37°C to an OD_600_ ∼ 0.4. Cells were then pelleted and resuspended in 30 ml of 25 mM Tris-HCl (pH 7.5), 100 mM NH_4_Cl, 15 mM MgCl_2_, 5 mM β-mercaptoethanol. Cells were harvested for RNA purification or flash frozen in liquid nitrogen and stored at −80°C.

### Chemical probing for m7G

Chemical probing for 7-methylguanosine in *E. coli* 16S rRNA was carried out essentially as described previously (Recht et al. 1996). Approximately 10 pmol 16S rRNA or RNA extracted from 70S ribosomes was resuspended in 20 µl 0.5 M Tris-HCl (pH 8.2). Selective reduction of m7G was performed by adding 5 µl freshly made 0.5 M sodium borohydride solution. The reaction was incubated on ice in the dark for 30 minutes. The reaction was ended by the addition of 10 µl 3 M sodium acetate (pH 5.2) and precipitated with ethanol. Pellets were washed once with 80% ethanol. RNA was pelleted by centrifugation and resuspended in 20 µl 1 M aniline/glacial acetic acid solution (1:1.5) (pH 4.5). RNA cleavage proceeded by incubating the reaction at 60°C for 10 minutes in the dark. The reaction was ended by the addition of 20 µl 0.5 M Tris-HCl (pH 8.2), and the RNA was isolated by extracting with phenol/chloroform/isoamyl alcohol. RNA was precipitated from the aqueous phase, pelleted and washed with 80% ethanol. RNA pellets were resuspended in 2.5 µl nuclease free water. Primer extension to determine the site of m7G-specific cleavage was carried out as described (Merryman and Noller 1998). To detect G527 methylation, the primer 5′-CGTGCGCTTTACGCCCA-3′ was used.

### MinION sequencing of 16S rRNA

MinION sequencing of 16S rRNA libraries was performed using MinKNOW version 1.1.30. The flow cells used were FLO-MIN106 SpotON version. ONT’s Metrichor base-calling software (1D RNA Basecalling for FLO-MIN106 v1.134 workflow) takes this raw signal and produces base-called FASTQ sequence in the 5′ to 3′ order after reads are reversed. During the course of these experiments, ONT made a new local base-caller available, named Albacore. We performed base-calling for the sequencing runs using Albacore v1.0.1, and performed all alignment-based analyses with the newer sequence data.

### Data analysis

FastQ sequences were extracted using poretools v0.6 ^32^ and then sequence alignment was performed using marginAlign v0.1 ^16^ (using BWA-MEM version 0.7.12-41044; parameter “-x ont2d” ^33^). The statistics were calculated using marginStats v0.1 ^16^. We then created assembly hubs to visualize these alignment on the UCSC genome browser using createAssemblyHub utility in marginAlign suite ^16^. We calculated read identity as matches / (matches + mismatches + insertions + deletions). We used marginAlign expectation maximization (EM) to estimate the error model from the sequence data. Using these high-quality alignments, we estimated substitution rates for the RNA nucleotides in MinION data. Using these high-quality alignments, we then performed variant calling using marginCaller v0.1 ^16^ to predict variants and associate systematic sequence mis-calls with putative base modifications. To test for systematic k-mer biases in MinION RNA data, we compared 5-mers in reads and the known 16S rRNA reference.

### Nanopore ionic current visualization

We used nanoraw v0.4.2 ^34^ to visualize ionic current traces for 16S rRNA reads from different *E. coli* strains that were sequenced on the MinION. We used the software with its default settings. We chose graphmap ^35^ as the aligner in nanoraw, and the argument ‘ont’ (now ‘pA’ in nanoraw v0.4.2) as the option for normalizing raw ionic currents. The ionic current plots were created using the plot_genome_location function. For all of the ionic current analysis, we inverted the reference sequence since the present MinION direct RNA sequencing chemistry sequences native RNA molecules in the 3′-5′ direction.

### Microbial classification

Binning reads by length (200-600, 600-1000, >1000 bases), we randomly sampled 10,000 reads per bin for each microbe. These reads were then mixed *in silico* and aligned using marginAlign v0.1 ^16^. A read was called as correctly classified if it aligned to one of the 16S rRNA reference sequences for that microbe. 10 classification iterations were performed for each of the bins.

---

**Supplemental Fig. S1.**
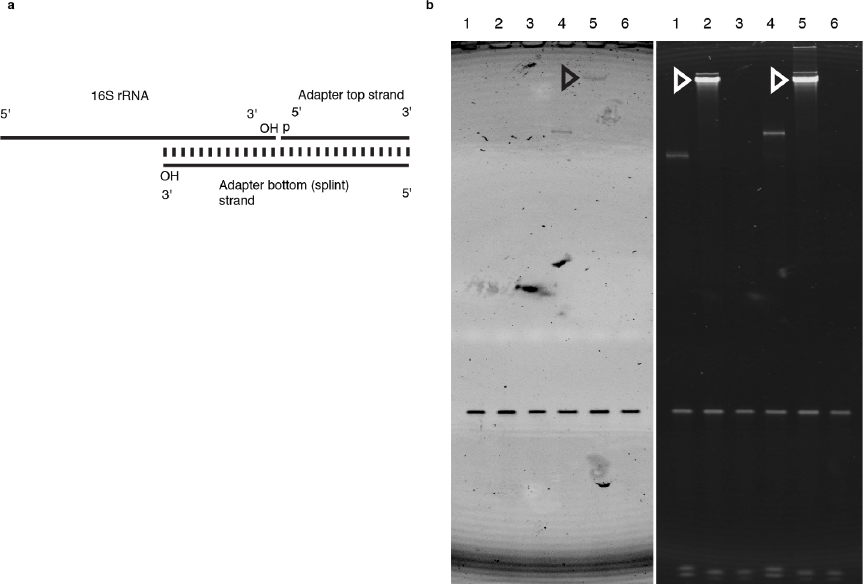
Design and testing of an oligonucleotide adapter to prepare 16S rRNA for nanopore direct RNA strand sequencing. **(a)** Schematic of oligonucleotide adapter hybridized to the 3′ end of 16S rRNA. The adapter bottom (splint) strand can hybridize to the conserved Shine-Dalgarno sequence on the 16S rRNA 3′ end. The adapter top strand can hybridize to the 5′ end of the adapter bottom strand as shown. The 3′ terminal hydroxyl of the 16S rRNA and the 5′ terminal phosphate of the adapter top strand can be covalently joined by T4 DNA ligase. **(b)** Denaturing gel analysis of a ligation reaction demonstrating the 16S adapter hybridizes and ligates to *E. coli* 16S rRNA 3′ ends. The left panel shows the unstained gel image. The lower band is a fluorescent, 3′-6-FAM-labeled version of adapter top strand. Lanes 1-3 show pre-ligation reaction samples for: Lane 1) negative control with just adapter present. Lane 2) positive control with an polyA-specific adapter containing a 3′ terminal oligo dT_10_ overhang (replaces 16S-specific overhang) and the 6-FAM labeled top strand. A synthetic 288mer polyA RNA is used as the control substrate. Lane 3) 6-FAM-labeled 16S rRNA-specific adapter and purified 16S rRNA from *E. coli*. Lanes 4-6 show post-ligation reaction samples for: Lane 4) negative control. Lane 5) positive control with polyA RNA 288mer. Lane 6) 16S rRNA reaction with 6-FAM labeled 16S rRNA-specific adapter. The size-shifted fluorescent top strand indicates ligation to the 16S rRNA 3′ end (Open arrow). The right image is the same gel stained with SybrGold. Position of the 16S rRNA is indicated by open arrows.

**Supplemental Fig. S2.**
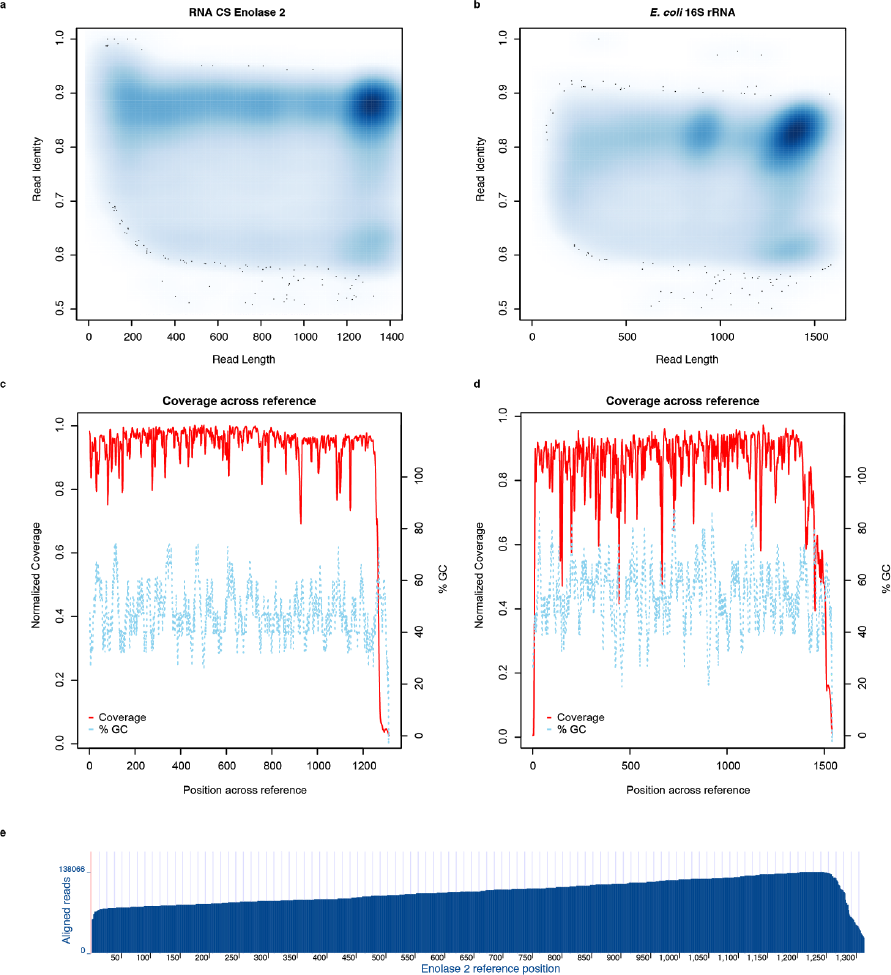
Alignment metrics for Enolase 2 polyA calibration strand and *E. coli* 16S rRNA. Alignments were performed using marginAlign (guide alignments from BWA MEM “-x ont2d” followed by chaining). (**a**) Identity vs. read length for Enolase 2. (**b**) Identity vs. read length for 16S *E. coli* rRNA. (**c**) Coverage across reference for Enolase 2 calibration strand. (**d**) Coverage across reference for 16S *E. coli* rRNA. (**e**) Alignment of 100,000+ Enolase 2 reads to the reference sequence.

**Supplemental Fig. S3.**
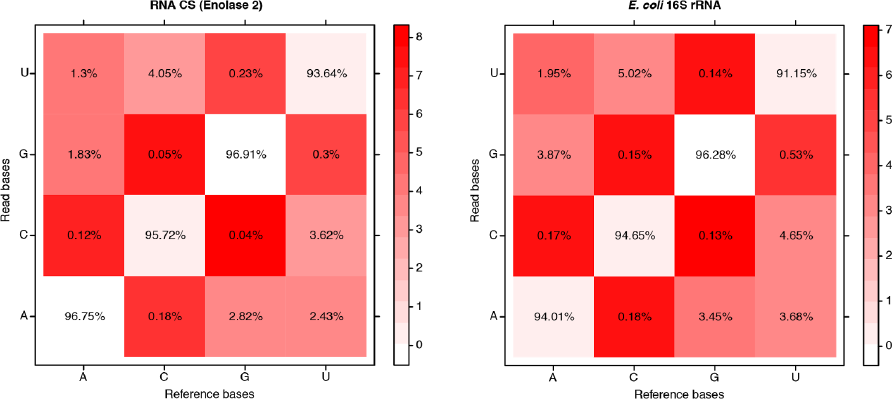
The matrix for substitution emissions for Enolase 2 calibration strand and *E. coli* 16S rRNA. This matrix was determined using marginAlign EM. The matrix shows low rates of C-to-G and G-to-C substitutions, relative to the other substitutions. The color scheme is fitted on a log scale, and the substitution values are on an absolute scale.

**Supplemental Fig. S4.**
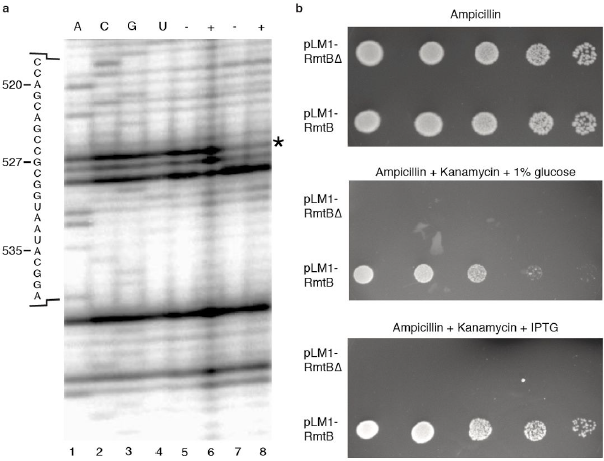
Confirmation of guanosine N7-methylation (m7G) at positions 527 and 1405 in E. coli 16S rRNA. **(a)** Canonical m7G527 is present in wild type *E. coli* and absent in the RsmG deficient *E. coli* strain. Sodium borohydride/aniline cleavage was used to detect N7-methylated guanosine in 16S rRNA for *E. coli* str. MRE600 (wild type) bearing m7G527 and RsmG deficient (mutant) *E. coli* str. BW25113 JW3718Δ. Lanes 1-4 are sequencing lanes for A, C, G, and U respectively. Wild type 16S rRNA from *E. coli* str. MRE600 is used as the template. Lanes 5 and 7: sodium borohydride/aniline treatment (labeled +) of 16S rRNA from wild type *E. coli* and 16S rRNA from RsmG mutant *E. coli*, respectively. Strand cleavage should result in an primer extension stop 1-nt ahead of G527 (position 527 marked by an asterisk). Lane 6 and 8: untreated 16S rRNA for wild type and mutant 16S rRNA. Primer extension products were run on denaturing 6% acrylamide gel, and imaged using a phosphorimager. **(b)** RmtB confers a kanamycin resistance phenotype consistent with G1405 N7-methylation in 16S rRNA from an engineered *E. coli* strain. Serial dilutions from 10^-2^ to 10^-6^ (Left to Right) of *E. coli* BL21 DE3 pLysS strains transformed with pLM1-RmtB and negative control pLM1-RmtBΔ were spotted on LB agar plates. The pLM1 plasmids use pET32a as the backbone, which contains an ampicillin resistance gene. The RmtB gene is under the control of a lactose inducible T7 promoter. Plates are supplemented with: 100 µg/ml Ampicillin (top), 100 µg/ml Ampicillin + 200 µg/ml Kanamycin + 1% glucose (middle), 100 µg/ml Ampicillin + 200 µg/ml Kanamycin + 1 mM IPTG (bottom).

**Supplemental Fig. S5.**
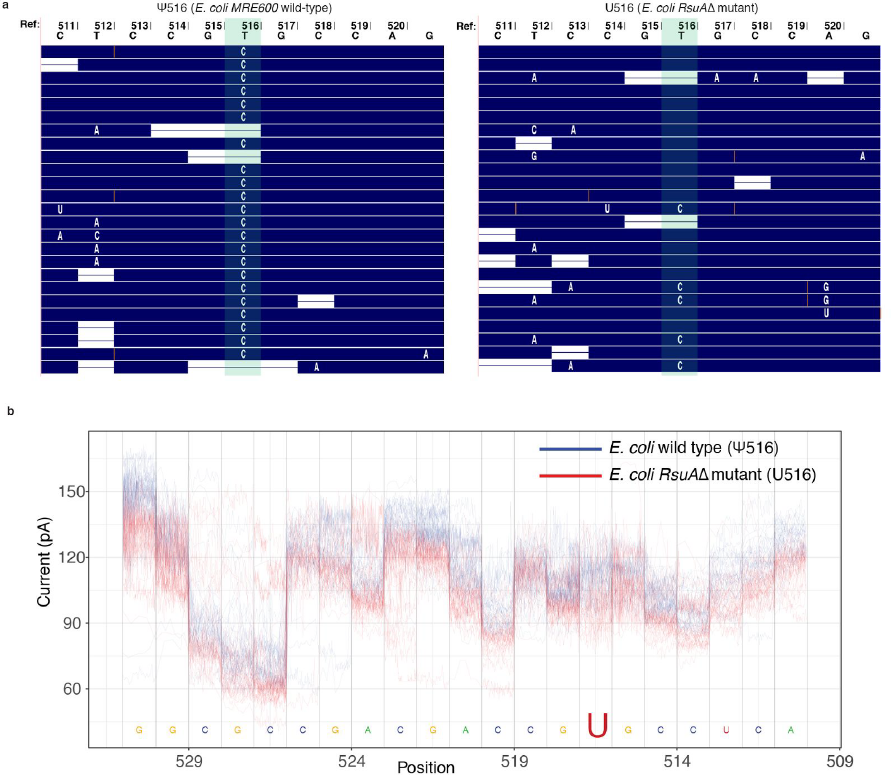
Inference of pseudouridine in *E. coli* 16S rRNA direct sequencing reads. **(a)** Comparison of aligned reads from strands containing putative pseudouridine versus strands bearing canonical uridine at position 516. Reads are aligned to the *E. coli* MRE600 *rrnD* 16S rRNA reference sequence. Shown are twenty-five 16S rRNA reads from separate sequencing runs for *E. coli* str. MRE600 (wild type), which bears a pseudouridine at U516 (ψ516) and an *RsuA* deficient strain (*RsuA*Δ mutant), which has a canonical U at position 516. Green shading indicates the position of U516 (shown as a T in the reference gene sequence). **(b)** Aligned ionic current traces from approximately thirty 16S rRNA reads covering position U516 from wild-type *E. coli* and *RsuA*Δ mutan strain. Pseudouridylation site, U516, is shown in large font. The sequence is shown 3′-to-5′ because ionic current signal is 3′-to-5′. Numbering uses standard *E. coli* 16S rRNA numbering.

**Supplemental Fig. S6.**
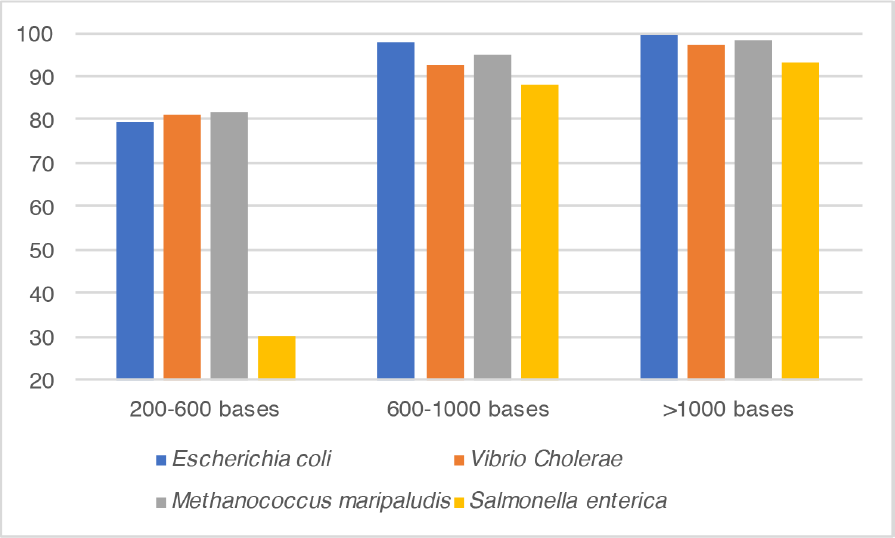
Microbial classification accuracy (%) using all MinION data vs. 16S rRNA read length. The classification was performed from an *in silico* mixture of 16S rRNA reads from four microbes. Reads were binned based on length. A read was called as correctly classified if it aligned to one of the 16S rRNA reference sequences for that microbe.

**Supplemental Table S1.** Error rate profile for Enolase 2 transcript and 16S *E. coli* rRNA. Error models were estimated using marginAlign (guide alignments from BWA MEM “-x ont2d” followed by chaining). Statistics were generated using marginStats.

| **Median** | **RNA CS (Yeast Enolase 2; 1.35 kb)** | **16S *E. coli* rRNA (1.6 kb)** |
| --- | --- | --- |
| **Alignment identity** | 87.10% | 81.59% |
| **Insertions** | 2.69% | 1.90% |
| **Deletions** | 2.97% | 6.02% |
| **Mismatches** | 5.20% | 7.20% |
| **Read coverage** | 96.14% | 97.57% |
| **Read length** | 1256 bases | 1349 bases |

**Supplemental Table S2.**
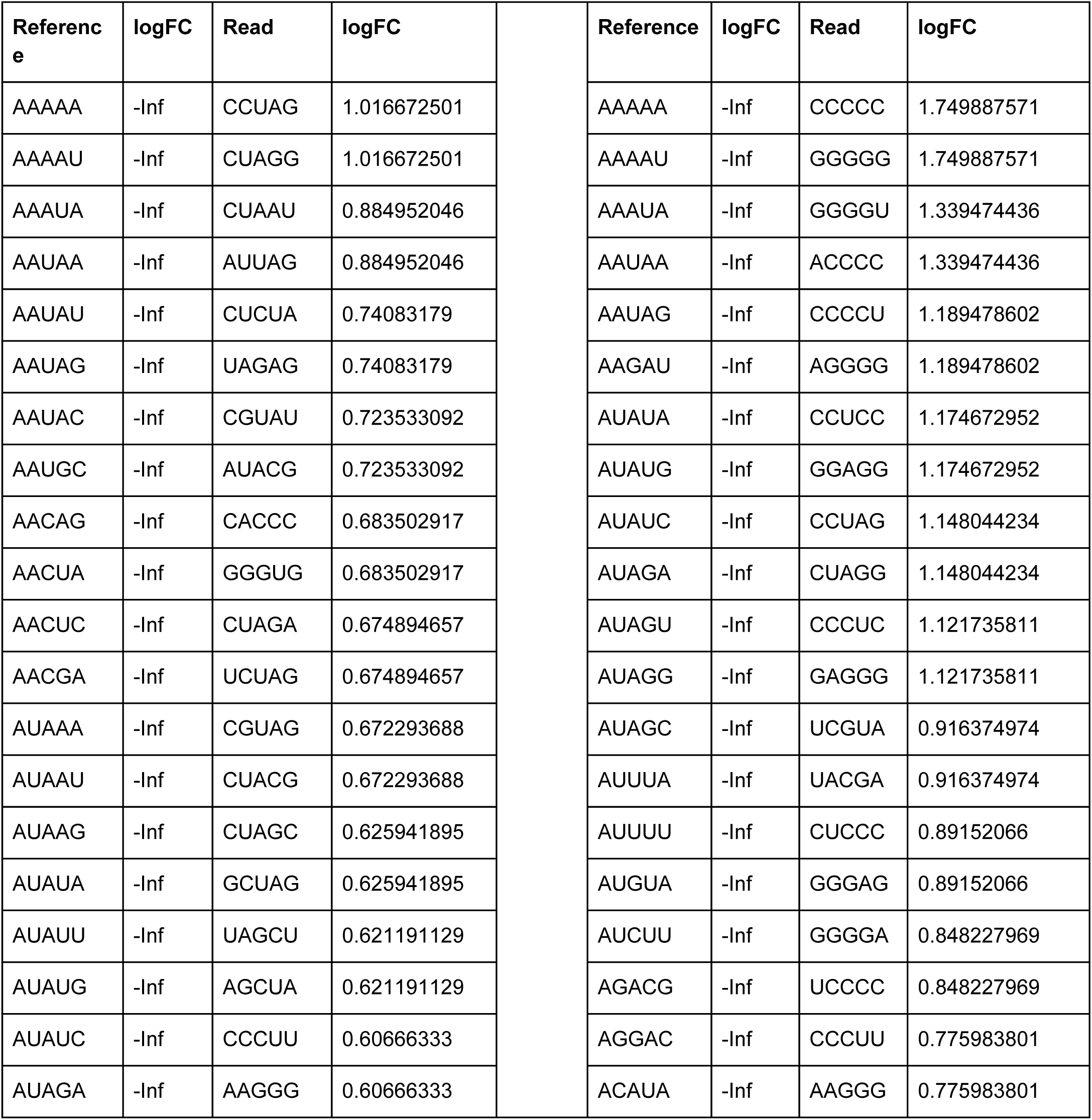
Over and Under represented 5-mers comparison for Enolase 2 (left) and *E. coli* 16S rRNA (right). 5-mers were counted and compared for RNA read data and their respective reference sequences. LogFC represents log fold-change.

**Supplemental Table S3.** Detection of nucleotide variants in 16S *E. coli* rRNA using marginCaller.

| ##fileformat=VCFv4.2 |  |  |  |  |  |  |  |
| --- | --- | --- | --- | --- | --- | --- | --- |
| ##fileDate=20170416 |  |  |  |  |  |  |  |
| ##source=marginCaller |  |  |  |  |  |  |  |
| ##reference=Ec_albacore_output/processedReferenceFastaFiles/ec_16S.fa |  |  |  |  |  |  |  |
| #CHROM | POS | ID | REF | ALT | QUAL | FILTER | INFO |
| ecoli_MRE600 | 6 | . | G | A | . | PASS | 0.323009035 |
| ecoli_MRE600 | 79 | . | A | G | . | PASS | 0.477024303 |
| ecoli_MRE600 | 90 | . | U | C | . | PASS | 0.495124202 |
| ecoli_MRE600 | 183 | . | C | U | . | PASS | 0.334402802 |
| ecoli_MRE600 | 226 | . | G | A | . | PASS | 0.455808427 |
| ecoli_MRE600 | 273 | . | U | A | . | PASS | 0.304643919 |
| ecoli_MRE600 | 288 | . | A | G | . | PASS | 0.38459441 |
| ecoli_MRE600 | 328 | . | C | U | . | PASS | 0.340478205 |
| ecoli_MRE600 | 346 | . | G | A | . | PASS | 0.301363107 |
| ecoli_MRE600 | 485 | . | U | C | . | PASS | 0.760495972 |
| ecoli_MRE600 | 516 | . | U | C | . | PASS | 0.908639215 |
| ecoli_MRE600 | 527 | . | G | C | . | PASS | 0.778486859 |
| ecoli_MRE600 | 790 | . | A | G | . | PASS | 0.496515677 |
| ecoli_MRE600 | 893 | . | C | U | . | PASS | 0.43893419 |
| ecoli_MRE600 | 1150 | . | A | U | . | PASS | 0.512479558 |
| ecoli_MRE600 | 1195 | . | C | U | . | PASS | 0.701573199 |
| ecoli_MRE600 | 1281 | . | U | C | . | PASS | 0.702653464 |
| ecoli_MRE600 | 1304 | . | G | A | . | PASS | 0.605698867 |
| ecoli_MRE600 | 1380 | . | U | C | . | PASS | 0.40811185 |
| ecoli_MRE600 | 1406 | . | U | C | . | PASS | 0.326475023 |
| ecoli_MRE600 | 1421 | . | G | A | . | PASS | 0.317460212 |
| ecoli_MRE600 | 1495 | . | U | A | . | PASS | 0.568537937 |
| ecoli_MRE600 | 1518 | . | A | U | . | PASS | 0.390200124 |
| ecoli_MRE600 | 1519 | . | A | U | . | PASS | 0.467437353 |

